# Epigenetic regulation clocks the multigenerational olfactory imprinting in *C. elegans*

**DOI:** 10.1101/2021.06.30.449753

**Authors:** Madeleine Erard-Garcia, Diana Andrea Fernandes de Abreu, Antoine Gruet, Marie-Pierre Blanchard, Kevin Baranger, Yoanne Clovis, François Féron, Jean-Jacques Remy

## Abstract

Imprinting is an early sensory life experience that induces adult behaviours, such as mother recognition or homing. In a previous study, we demonstrated a striking olfactory imprinting in *C. elegans* that can be inherited over generations. When exposed to specific odorants during a timely controlled post-hatch period, *C. elegans* worms display during adulthood an enhanced migration towards these molecules. In order to unveil some of the genetic and epigenetic factors that are responsible for such a behavioural plasticity, we assessed the role of heterochronic genes using a candidate gene approach. We report here that translation of the Hunchback-Like 1 (HBL1) transcription factor in the sensory processing interneuron AIY, is a determining factor for olfactory plasticity timing in *C*.*elegans*. HBL1 may associate to the SPR1/CoREST co-repressor, the lysine demethylase SPR5/LSD1 and the histone deacetylase HDA3 to lengthen the plasticity period, whereas the translation initiation factor IFE-4 and the histone deacetylase HDA2 abridge it. We also observed that lengthened plasticity periods allow proportionally faster stable behavioral adaptation of *C. elegans* populations. We conclude that plasticity timing is a key factor, not only to transiently adapt individuals but also to stably adapt animal populations via multigenerational accumulation of experience.

## Introduction

Behavioural adaptation is determined by a continuous intertwining of genetic and experiential factors, in particular during critical periods of plasticity. During these early life time-windows, juvenile animals receive and integrate environmental information that will imprint their adult behaviour.

Behaviour and cognitive abilities in birds are the best-described examples of how critical periods of plasticity are linked to behavioural adaptation (1). Bird species are classified along a spectrum spanning from precocial to altricial, with established differences in brain size at hatching, timing of brain development and behavioural complexity. Precocial birds, as Megapodes and Gallinaceous, rapidly express adult innate behaviours and become fully autonomous within hours or days post-hatching. In contrast, adult behaviours are expressed more slowly in altricial birds (parrots, passeriforms and song birds) that need up to weeks-long critical periods of nurturing, from environmental stimuli and parental care, to fully express adult behaviours. Evolution within the precocial/altricial spectrum is not limited to the bird class: growing complexity of cognitive capacities has also been linked to lengthening juvenile periods of plasticity, during primate (2) and hominid (3) evolution.

The *C. elegans* animal model is innately attracted to a number of olfactory cues, such as benzaldehyde (BA) and citronellol (CI) (4). These odorants bind the olfactory neuron receptors AWC (4) and the olfactory information is further processed by post-synaptic interneurons AIY. We show here that chemo-attraction is significantly reduced when induced during the course of the juvenile period, in comparison with the later stages of development. *C*.*elegans* worms however exhibit a critical period for chemosensory behaviour plasticity, which takes place during the first 12 hours after hatching (5, 6).

We previously demonstrated that an exposition of juvenile animals to BA or CI during this critical period of plasticity induces a life-long behavioural adaptation via olfactory imprinting. At the adult stage, odour-exposed animals display an enhanced migration towards BA or CI (chemo-attraction) when compared to naïve animals (5). Strikingly, we also observed that olfactory imprinting is inherited either transiently or stably, depending on the number of consecutive worm generations exposed to BA or to CI during the period of olfactory plasticity. In the wild-type N2 strain of *C*.*elegans*, stable trans-generational inheritance of olfactory imprinting is precisely observed after five consecutive generations were odour-exposed (7).

The chemosensory behaviour is thus timely regulated in *C. elegans* worms. Since no behavioural timer genes have been identified in the worm so far, we used a candidate gene approach.

A well-described developmental heterochronic pathway controls the succession of the four L1 to L4 larval stages preceding the development of adult egg-laying worms. Mutations in the developmental heterochronic genes can disrupt the right succession of developmental stages by producing premature or delayed terminal differentiation phenotypes (8-10). We first considered the zinc finger transcription factor Hunchback-like-1 (HBL-1) as a chemo-attraction timer, as HBL-1 controls the synchrony of several developmental events in *C. elegans* (11-15). In addition to be part of the developmental heterochonic pathway, HBL-1 is also the worm ortholog of the *Drosophila melanogaster* Hunchback transcription factor (*hb*), a well-known timer of neuronal maturation in the fly (16). d4EHP, an eIF4E-like cap-binding protein, was shown to repress *hb* while establishing axis polarity in the early drosophila embryo (17). Strikingly, one of the worm eIF4E translation initiation factor, IFE-4, was also reported to affect the translational efficiency of *hbl-1* mRNA in *C. elegans* (18).

When compared to the wild-type N2 worms, the juvenile *hbl-1* mutants worms precociously displayed chemo-attraction while they lost olfactory plasticity and were unable to imprint odours. By contrast, the developmental expression of chemo-attraction was strongly delayed in the *ife-4* mutants worms, compared to N2. In addition, the *ife-4* worms keep a life-long ability to imprint odours, up to five times longer than wild-type animals. Moreover, the *ife-4* mutation maintained high levels of the HBL-1 protein all over the development, suggesting the timing of olfactory plasticity and imprinting results from a translational regulation of *hbl-1* by the translation initiator IFE-4.

We further reasoned that mutations in genes that control the timing of neuronal maturation would also affect the whole temporal settings of chemo-attractive responses. Neuronal timers in other species, as *hb* and Tramtrack88 (Ttk88) in *Drosophila* (16, 19) and REST/NRSF in mammals (20, 21), are known to epigenetically control neuronal gene expression through association with the CoREST co-repressor and with conserved chromatin modifying factors as the Lysine-specific demethylase-1 (LSD-1) and the Class I Histone deacetylases (HDAC) (22). We report here that genetic inactivation of SPR-1, the worm ortholog of CoREST (23), or of the chromatin modifying factors SPR-5, HDA-2 and HDA-3, respectively the worm orthologs of the mammalian LSD1, HDAC-2 and HDAC-1, desynchronized the worm chemosensory behaviour.

We found that all mutations that extended the imprinting period in individuals proportionally decreased the number of generations to be imprinted before a stable change in chemosensory responses in populations. From these observations, we inferred that a stable behavioural change might be triggered after a fixed amount of heritable olfactory experience has been cumulated through generations.

## Results

### Biphasic implementation of chemo-attraction behaviour throughout *C. elegans* development

*C. elegans* is innately attracted to BA and CI, with maximal attraction to the 1/100 dilution. The life cycle of *C. elegans* is comprised of the *in ovo* embryonic stage, four larval stages and adulthood starts at 60 hours post-hatching.

We evaluated innate BA chemo-attraction at different time-points of N2 wild-type *C. elegans* development in order to assess its implementation (for a detailed description see Materials and Methods). BA chemo-attraction implemented throughout development (Figure 1). From 10 hours post-hatching, chemo-attraction progressed with a slow rate of implementation (flat slope of 0.015) while it progressed with a higher rate of implementation from 38 hours post-hatch to adulthood (steeper slope of 0.07), reaching its maximum level at the age of 96 hours. The same biphasic progression was observed for the implementation of chemo-attraction to CI (data not shown), sensed by the same AWC olfactory neuron (24). This biphasic implementation contrasts with the almost linear size progression of the *C. elegans* larvae (25).

**Figure 1.**
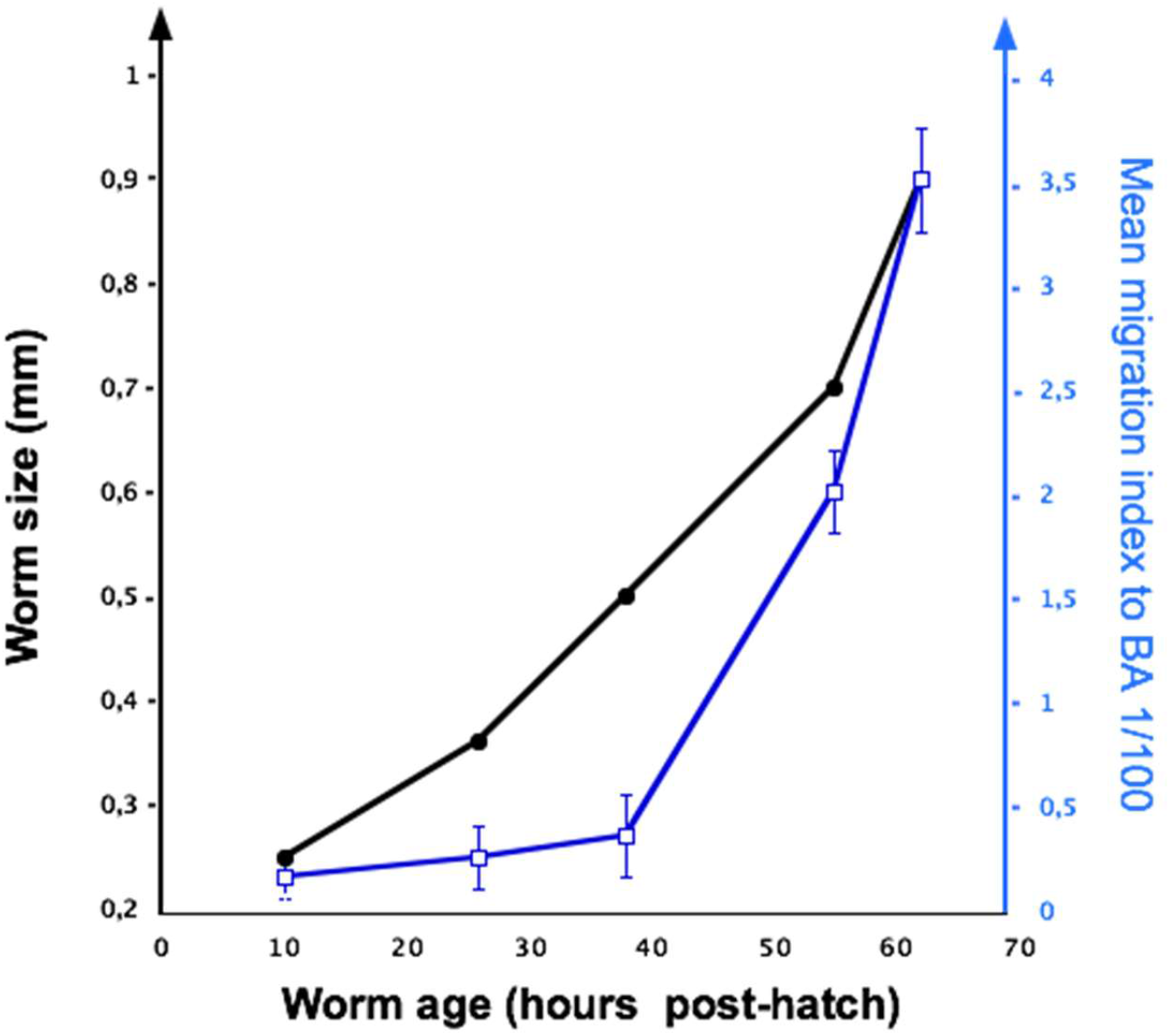
Implementation of chemo-attraction during physiological *C*.*elegans* development. BA 1/100 chemo-attraction of wild-type N2 worms was measured at 10, 26, 38, 56 and 60 hours post-hatch (blue curve). The mean migration indices (MMI) were calculated as indicated in Supplemental Material and Methods. Each chemotaxis assay included 20 worms and was repeated four times. The black curve depicts the wild-type growth curve of *C. elegans* (as described in WormAtlas, Altun, Z. F. and Hall, D. H. (ed.s). 2002-2006).

### Hunchback-like-1 (HBL-1) may act as a timer of chemo-attraction implementation during the course of development

To identify potential molecular timers, the early innate BA chemo-attraction implementation was assessed in larvae with mutations in *hbl-1*, in *ife-4*, and in worms with a transgene over-expressing *hbl-1* solely in AIY. The 10, 15 and 26 hours old *hbl-1 (ve18)* and *hbl-1 (mg285)* mutant larvae migrate in BA gradients significantly faster than N2 wild-type larvae of same age (Figure 2A). 10 hours old *hbl-1* larvae migrate as fast as 26 hours old N2 larvae, suggesting the absence of a functional *hbl-1* gene hastened the onset of BA chemo-attraction by at least 16 hours. In contrast, the innate chemo-attraction to BA was significantly delayed in *ife-4 (ok320)* mutants and in *ttx3::hbl-1* transgenic larvae, compared to N2. 26 hours old *ife-4* and *ttx3::hbl-1* larvae behaved as 10 hours old N2 larvae, indicating a mean delay of 16 hours.

**Figure 2.**
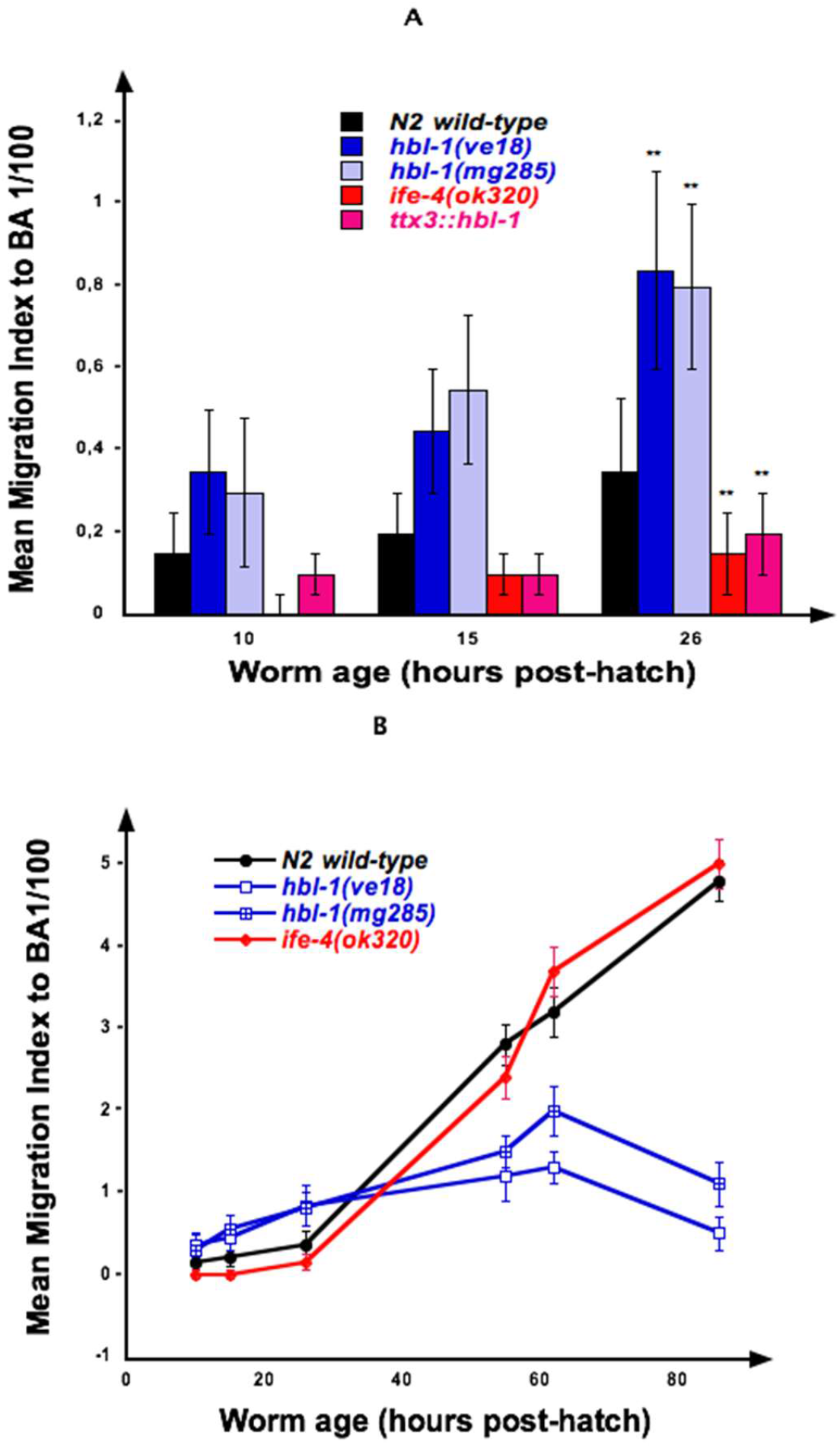
Implementation of chemo-attraction depends on HBL-1. **A. Implementation of BA-chemo-attraction during early life in *C***.***elegans* mutants**. Chemotaxis assays were performed at age 10, 15 and 26 hours post-hatch in N2, in *hbl-1* (*ve18* and *mg285), ife-4 (ok320) mutants* and in *ttx-3::hbl-1* transgenic. Mean migration indices were calculated for each chemotaxis assay and compared to N2. Each chemotaxis assay included 20 worms and was repeated four times. **p-value < 0.01. **B. Implementation of chemo-attraction throughout development in *C***.***elegans* mutants**. BA 1/100 chemo-attraction of wild-type N2, hbl-1, ife-4, and ttx-3::hbl-1 mutants was measured at 10, 15, 26, 56, 60 and 84 hours post-hatch. Mean migration indices were calculated for each chemotaxis assay and compared to N2. Each chemotaxis assay included 20 worms and was repeated four times.

We further compared the innate BA chemo-attraction in *hbl-1, ife-4* and N2 worms during the whole course of development until the adult stage (Figure 2B). At early stages, BA attraction progressed two times faster in *hbl-1* mutants than in N2, with progression slopes of 0.031 and 0.015, respectively. Impaired motility at later stages in *hbl-1* mutants is due to well-described morphological defects (12). The mean migration of *ife-4 (ok320)* mutants in BA chemo-attraction tests becomes similar to N2 after 26 hours post-hatch, implementing with the same rate (slope of 0.07).

These observations prompted us to investigate *hbl-1* and *ife-4* transcription levels in the AIY interneuron of wild-type N2 using fluorescent reporters (Supplemental Figure 1 A and B). To assess *hbl-1* expression, we analyzed a GFP reporter construct containing the *hbl-1* promoter and 1 kb of the *hbl-1* gene (reporter strain BW1932 (12)). To examine the pattern of *ife-4* expression we used the transgenic extra-chromosomal array *ife-4*::GFP(*sEx15051)*. In addition, a reporter construct expressing CherryFP under the AIY specific *ttx-3* promoter B was used to label the AIY interneuron (strain ott4607; kind gift from O. Hobert).

We observed the expression of the *hbl-1::gfp-hbl-1* reporter overlaps with the AIY specific label at all stages (Supplemental Figure 1A), suggesting *hbl-1* transcription in AIY might not be down-regulated after the early olfactory plasticity and imprinting period. Interestingly, *ife-4::GFP* is detected in AIY but not before 12 hours post-hatch, which corresponds to the end of the imprinting period. The *ife-4*::GFP reporter is also expressed in other neuronal cell bodies (Supplemental Figure 1B).

We therefore hypothesized that IFE-4 protein could inhibit HBL-1 protein synthesis ending the period of plasticity. In order to test this, we quantified the levels of HBL-1 in ife-4, in hbl-1 mutants and in wild-type N2 worms at several developmental stages, using home made HBL-1 specific antibodies (Supplemental Figure 2). In N2, the level of HBL-1 in AIY might be higher at the earlier than at the later larval stages. HBL-1 in N2 larvae was however only detected after loading high amount of larval protein extracts (Supplementary Material and Methods). This technical limitation probably explains why a down-regulation of HBL-1 is not observed in N2.

The *ife-4 (ok320)* mutants synthesised higher levels of HBL-1 protein than N2 at all developmental stages, indicating IFE-4 is indeed acting as a translational inhibitor of *hbl-1* (Supplemental Figure 2).

Compared to wild-type, chemo-attraction is precociously expressed in *hbl-1* mutants, while over-expression and over-time persistence of HBL-1 in the *ife-4* mutants and in the *ttx-3::hbl-1* transgenic animals delayed chemo-attraction expression. Taken together this body of data strongly suggests that *hbl-1* translation in the AIY neuron is a main timing factor for chemo-attraction expression during early development.

### Plasticity time-length is negatively correlated to chemo-attraction expression

We hypothesised the expression of innate chemo-attraction might be put on hold during the olfactory plasticity period during which olfactory experience can be imprinted. If true, delayed expression of chemo-attraction would be linked to lengthened imprinting periods, while precocious expression of chemo-attraction would be linked to shortened imprinting periods.

In order to test this hypothesis, precocious *hbl-1* and delayed *ife-4* and *ttx-3::hbl-1* mutants were exposed to BA 1/300 during 24 hours post-hatch, a time interval that spans the wild-type imprinting period (Figure 3). The behaviour of BA-exposed and the behaviour of unexposed worms from each strain were then compared at four days post-hatch. Imprinting index is calculated by subtracting the responses of naive from the responses of odour-exposed worms (for a detailed description see Materials and Methods). A positive imprinting index thus indicates odour-exposed worms migrated faster than naïve-unexposed worms.

**Figure 3.**
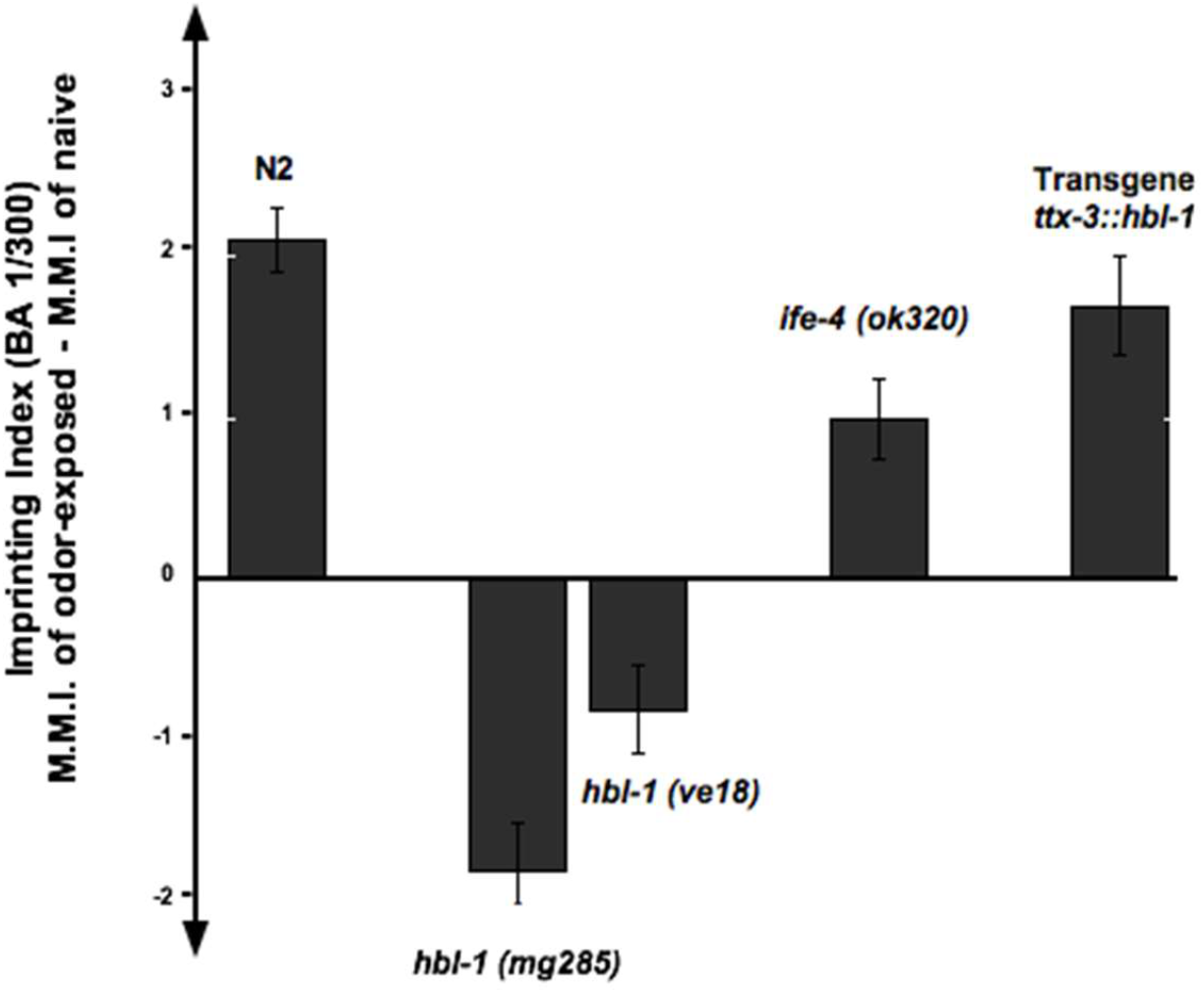
Olfactory imprinting is HBL-1 dependent. Wild-type N2 and hbl-1, ife-4 and ttx-3::hbl-1 mutants were exposed to BA 1/300 for 12 hours post-hatch in order to generate an olfactory imprinting. Chemotaxis assays of these odor-exposed and their non-exposed counterpart were then performed at adulthood. Mean imprinting indices to BA 1/300 were calculated by subtracting the MMI of non-exposed mutants from the MMI of odor-exposed mutants (detailed description in Material and Methods). Each chemotaxis assay included 20 worms and was repeated four times. **p-value < 0.01.

BA pre-exposure enhanced adult BA attraction in N2 wild-type worms, with a positive mean imprinting index of 2.1 ± 0.25. BA pre-exposure also enhanced adult BA chemo-attraction in the delayed *ife-4* and *ttx-3::hbl-1* mutant worms with positive imprinting indices of 1 ± 0.25 and 1.7 ± 0.3, respectively (Figure 3). Early BA exposure of the precocious *hbl-1 (ve18)* and *hbl-1 (mg285)* mutants induced by contrast a decreased chemo-attraction to BA. This led to negative mean imprinting indices of - 0.8 ± 0.3 for *hbl-1 (ve18)* and of - 1.8 ± 0.3 for *hbl-1 (mg285)* worms, respectively (Figure 3).

### Histone-modifying factors are behavioural plasticity timers

In order to identify other chemo-attraction timers, we considered the *REST (RE*1-*S*ilencing *T*ranscription Factor) pathway. REST is a mammalian timer of neuronal differentiation (21). It mediates transcriptional repression in association with its co-repressors mSin3 and CoREST. In addition, the REST-CoREST complex can recruit histone modifiers as histone deacetylases (HDACs) and histone demethylase (LSD1) for long-term silencing of neuronal genes (21, 22). In the worm, no REST ortholog has been characterized, but a core complex consisting of SPR-1 (CoREST ortholog), SPR-5 (LSD1/KDM1 ortholog) and, possibly yet unknown, HDACs, might be recruited to the regulatory region of target genes through interaction with unidentified transcription factors with REST function (23).

HBL-1 may have a REST-like activity in the *C. elegans* neurons, as Hunchback in *Drosophila melanogaster*. HBL-1 may thus associate with CoREST/SPR-1, LSD1/SPR-5 and with some HDACs to time the worm chemosensory behavior. If so, mutations in the corresponding genes would also modify the onset of chemo-attraction.

In order to test this hypothesis, the innate chemo-attraction of several mutants of this putative timing complex was assessed. Two *spr-1 (ar205)* or *spr-1 (ar200)* mutants, lacking the C-terminal SANT domain of SPR-1 (23), exhibited a precocious innate BA chemo-attraction at 26 hours post-hatch, similarly to hbl-1 mutants (Figure 4A). We assessed three spr-5 alleles *ar197, by101* and *by119*. Only *spr-5 (by119)* larvae exhibited a precocious innate chemo-attaction, compared to wild-type. This discrepancy may be due to the nature of the mutations. The structure of the putative worm SPR-1/SPR-5 complex is unknown. Based on the 3D structure of the mammalian CoREST/LSD-1 complex (26), the *ar197* and *by101* mutations may only affect the C-terminus of the SPR-5 protein, leaving intact the putative SPR-1 interacting domain. By contrast, the *by119* allele, carrying a premature stop codon in the FAD-binding domain, may impair the formation of a SPR-1/SPR-5 complex.

**Figure 4.**
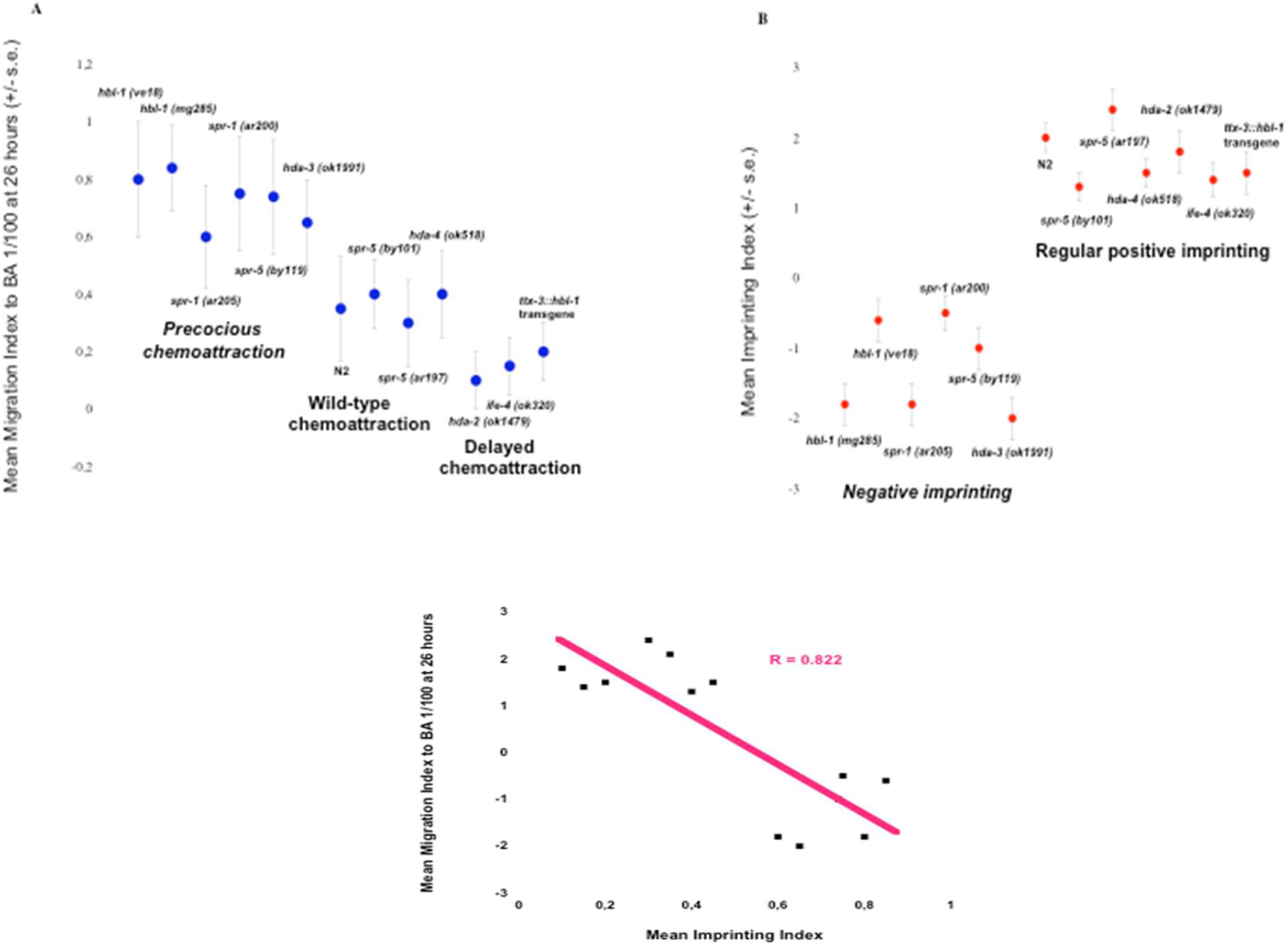
Precocious implementation of chemo-attraction is negatively correlated to chemo-attraction expression. **A. Implementation of chemotaxis response to BA** Mutant larvae aged of 26 hours post-hatch were assayed for chemo-attraction to BA 1/100. Mean migration indices were then calculated for each chemotaxis assay as described in Supplemental Material and Methods. Each chemotaxis assay included 20 worms and was repeated four times. **B. Acquisition of olfactory imprinting** Worms were exposed to BA 1/300 for 12 hours post-hatch in order to generate an olfactory imprinting. Chemotaxis assays of these odor-exposed and their non-exposed counterpart were then performed during adulthood. Mean imprinting indices to BA 1/300 were calculated by subtracting the MMI of non-exposed mutants from the MMI of odor-exposed mutants (detailed description in Material and Methods). Each chemotaxis assay included 20 worms and was repeated four times.

The *C. elegans* Class I HDAC is made of three genes, *hda-1, hda-2* and *hda-3. hda-1* is needed for the development of the nervous system and axon guidance (27). The *hda-1* mutants are sterile and severely uncoordinated, and were thus excluded from our screen. We thereafter compared the innate BA chemo-attraction timing of worms with deletions in *hda-2* and *hda-3*, the two remaining Class I HDAC genes, as well as in *hda-4*, a Class II HDAC gene involved in chemoreceptor gene expression (28). While the *hda-4 (ok518)* deletion had no effect on chemo-attraction timing, the *hda-2 (ok1479)* and *hda-3 (ok1991)* deletions produced heterochronic phenotypes. The *hda-2 (ok1479)* larvae behaved as the delayed *ife-4 (ok320)* larvae, while the *hda-3 (ok1991)* larvae precociously expressed BA chemo-attraction compared with wild-type N2 (Figure 4A).

### Chemo-attraction timing is linked to the processing of olfactory experience

The mutant worms we have assessed can be divided into three groups according to innate BA chemo-attraction phenotypes displayed at 26 hours (Figure 4A). A first group with wild-type BA chemo-attraction comprises *spr-5 (ar197), spr-5 (by101)* and *hda-4 (ok518)*. A second group includes *ife-4 (ok320), hda-2 (ok1479)* and the *ttx-3::hbl-1* transgenic worms that display a delayed innate BA chemo-attraction. A third group is made of the *hbl-1 (mg285), hbl-1 (ve18), spr-1 (ar205), spr-1 (ar200), spr-5 (by119)* and *hda-3 (ok1991)* mutants that display a precocious innate BA chemo-attraction. In order to test how chemo-attraction implementation affects the olfactory imprinting, mutants within the three groups were exposed to BA 1/300 during 12 hours post-hatch, the wild-type imprinting period (Figure 4B). In wild-type N2, BA-exposure enhances adult response to BA, compared to naïve N2 unexposed worms. All mutants belonging to the first and second groups imprinted BA as wild-type N2 worms. Strikingly, early BA exposure of mutants belonging to the third group decreased BA-chemo-attraction. Mean imprinting indices were found all negative for worms carrying the *spr-1 (ar205), spr-1 (ar200), spr-5 (by119)* and *hda-3 (ok1991)* mutations, which reflect a lower BA chemo-attraction for BA-experienced than for naïve worms. Worms with these mutations do keep a lifelong negative imprint of early experience. As shown in Figure 4 inset, our behavioural screen thus disclosed a strict correlation (R = 0.822) between the early onset of BA chemo-attraction (Mean Migration Index to BA 1/100 at 26 hours) and the way early BA experience is accounted by adults (Mean Imprinting Index).

### Delayed onset of chemo-attraction is associated with extended time-lengths of olfactory plasticity

Mutations that delayed the expression of innate BA chemo-attraction might also extend the BA imprinting plasticity period. To test this, *hda-4 (ok518), hda-3 (ok1991), hda-2 (ok1479), ife-4 (ok320)* and *ttx-3::hbl-1* mutants were BA-exposed during 0-12, 0-60, 12-36, or 36-60 hours post-hatch. As shown in Table 1, N2 and *hda-4 (ok518)* worms exclusively imprinted the BA-chemo-attraction when exposed during time periods encompassing the wild-type N2 critical period (time intervals 0 to 12 and 0 to 60 hours). Negative imprinting in *hda-3 (ok1991)* worms happened during the same time periods. In contrast, the competence for producing an imprint persisted far beyond the wild-type olfactory plasticity period in the *hda-2 (ok1479), ife-4 (ok320)* and *ttx-3::hbl-1* mutant worms. *hda-2 (ok1479)* kept on generating BA imprints up to the age of 36 hours (as indicated by the 12-36 hours period), displaying an imprinting plasticity period at least three times longer than N2. *ife-4 (ok320)* mutants and *ttx-3::hbl-1* transgenics imprinted at all stages, displaying an imprinting plasticity period at least five times longer than N2. Noteworthy, the larval development in *hda-2 (ok1479), ife-4 (ok320)* and in the *ttx-3::hbl-1* mutants was unaffected (data not shown), suggesting these worms imprint odours efficiently at any age, irrespective of the developmental stage.

**Table 1.** Mutations that affect the timing of olfactory imprinting critical periods. Worms of the indicated genotypes were either exposed or not exposed to BA 1/300 during the indicated periods of post-hatch development in order to generate an imprint. Each chemotaxis assay included 20 worms and the mean migration indices each are indicated (n=number of repeats). MMI for odor-exposed and for naïve unexposed worms were compared using the Student *t* unpaired data with unequal variance test. All ***italic bold numbers*** indicate that p-value<0.01.

| Worm strain | BA 1/300 exposure time (post-hatch hours) |  |  |  |  | Critical period of imprinting |
| --- | --- | --- | --- | --- | --- | --- |
|  | Unexposed | 0 to 12h | 0 to 60h | 12h to 36h | 36h to 60h |  |
|  | Mean migration index (+/- s.e.) |  |  |  |  |  |
| *ife-4(ok320)*<br>*n*=5 | 1.7±0.3 | 2.6±0.2 | 3.3±0.25 | 2.9±0.25 | 3.4±0.3 | > 60h<br>*post-hatch* |
| *ttx-3 ::hbl-1*<br>*n*=4 | 1.6±0.25 | 2.8±0.28 | *nd* | *nd* | 3.2±0.25 | > 60h<br>*post-hatch* |
| *hda-2(ok1479)*<br>*n*=5 | 1.6±0.3 | 3.2±0.3 | 3.5±0.3 | 3.3±0.25 | 1.4±0.3 | 36h<br>*post-hatch* |
| *N2 wild-type*<br>*n*=10 | 1.65±0.3 | 3.6±0.25 | 3.4±0.3 | 1.8±0.2 | 1.55±0.2 | 12h<br>*post-hatch* |
| *hda-4(ok518)*<br>*n*=7 | 1.8±0.3 | 3.4±0.3 | 3.5±0.35 | 2.1±0.3 | 2.0±0.2 | 12h<br>*post-hatch* |
| *hda-3(ok1991)*<br>*n*=7 | 1.55±0.25 | -0.4±0.2 | -0.3±0.18 | 1.4±0.3 | 1.7±0.2 | 12h<br>*Negative* |

### Cumulated multi-generational experience determines the rate at which behaviour is stably reshaped

Olfactory imprinting is always passed to the unexposed F1 generation, but it is not transmitted to the unexposed F2 generation in wild-type N2 worms (7). However, if five successive worm generations are odour-exposed, all the following unexposed generations (F5+1, +2, +N unexposed generations) inherit the imprint (7). The multi-generational olfactory-experience has stably reshaped the chemo-attractive behaviour in worm population.

In this paper, we have identified mutants with variable imprinting periods: 12 hours long for N2, *hda-4 (ok518)*, 36 hours long for *hda-2 (ok1479)* and at least 60 hours long for *ife-4 (ok320)* and *ttx-3::hbl-1*. Considering this unique body of data, we could test if, as suggested (29), the accumulation of multigenerational imprinted experience is responsible for stably reshaping the behaviour of an animal population.

To test this hypothesis, we first determined if variable lengths of plasticity periods affected the rate at which BA-imprinting was stably assimilated by the progeny. Previous study showed that if imprinting is still present at the second, a fortiori at the fifth unexposed generation shown in Table 2, it will be permanently fixed and inherited by all following generations in worm populations (7). In order to assess this question, different mutants were BA-exposed for 60 hours post-hatch for five consecutive generations. Imprinted BA-exposition was stably inherited by the N2 or *hda-4 (ok518)* after five generations (Ngen = 5) had been BA-exposed but not before (Table 2). Strikingly, in the case of mutants with lengthened plasticity periods, the stable assimilation of BA-imprinting needed less generations than for N2 to happen. For the *hda-2* mutant, that has a 36 hours plasticity period, the stable assimilation of BA-imprinting happened after four generations of odour-exposure (Ngen = 4). Moreover, for *ife-4 (ok320)* and *ttx-3::hbl-1* mutants, that have a plasticity period of at least 60 hours, the assimilation of the BA-imprints happened immediately at the first generation (Ngen = 1). Figure 5A summarizes how the imprinting plasticity period influences the number of generations required before imprinting assimilation operates. These results strongly suggest that a total of 60 hours of olfactory imprinting during the plasticity period will reshape the innate behaviour of worm populations. In the case of wild-type N2 worms, one generation of imprinted BA-exposition would produce 1/5 (around 12 hours) of the amount of imprints required for behavioural reshaping. In order to test this hypothesis, consecutive generations of *ife-4 (ok320)* mutants were BA-exposed either during 12, 36 or 60 hours post-hatch per generation, corresponding respectively to the N2, to the *hda-2 (ok1479)*, and to the whole *ife-4 (ok320)* imprinting plasticity periods (Figure 5B). It took four generations with multigenerational 36 hours long exposures, and five generations with multigenerational 12 hours long exposures before stabilisation of the behaviour. As already shown, in *ife-4* mutants able to imprint during at least 60 hours, a single exposure of 60 hours led to stable inheritance. We inferred that at least 60 hours of imprinted olfactory experience must be accumulated to reach the amount of olfactory experience required for a stable behavioural change.

**Table 2.** Prolonged imprinting periods accelerate behavioural reshaping in worm populations. Worm mutants were exposed to BA 1/300 for 60 hours post-hatch in order to generate an olfactory imprinting. This odor-exposure was reiterated for 1, 4 or 5 consecutive generations (Ngen=1, Ngen=4 or Ngen=5). To assess stable reshaping of the BA-chemoattraction, the F1 to F5 progeny issued from each BA-exposed generation (and from unexposed as control) were grown without BA exposure for 5 generations. Mean migration indices of the unexposed progeny - from F1 up to F5 - were determined and compared to the mean migration indices of naïve animals of the same genotype, as described. Each chemotaxis assay included 20 worms and was repeated four times. All italic ***bold numbers*** indicate that p-value<0.01.

| Worm strain | BA 1/300 responses of the fifth unexposed generation after N generations (Ngen) were odor-exposed (0-60h) |  |  | Generation from which imprinting switch to stable |
| --- | --- | --- | --- | --- |
|  | Ngen=1 | Ngen=4 | Ngen=5 |  |
| *ife-4(ok320)* | *3.6±0.2* | *3.4±0.2* | *3.6±0.3* | **FIRST** |
| *Transgenics ttx-3 ::hbl-1* | *3.2±0.3* | *3.0±0.2* | *3.8±0.3* | **FIRST** |
| *hda-2(ok1479)* | *1.5±0.3* | *3.0±0.3* | *3.2±0.3* | **FOURTH** |
| *N2 wild-type* | *1.5±0.3* | *1.8±0.25* | *3.2±0.3* | **FIFTH** |
| *hda-4(ok518)* | *1.7±0.3* | *1.8±0.3* | *3.3±0.35* | **FIFTH** |
| *hda-3(ok1991)* | *-0.5±0.2* | *-0.25±0.3* | *-0.4±0.2* | **FIRST** |

**Figure 5.**
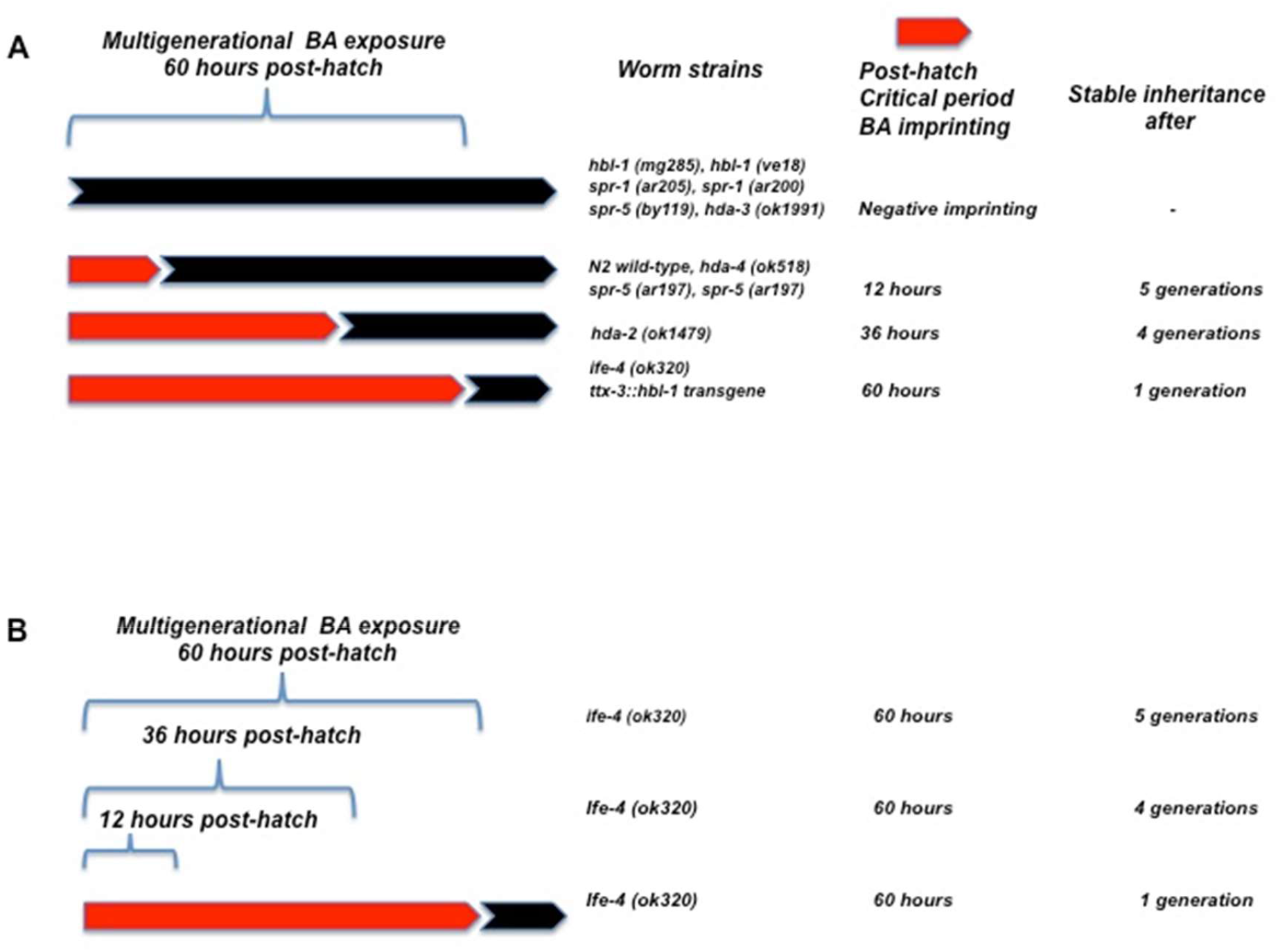
The rate of chemo-attraction behaviour reshaping is determined by the multigenerational accumulation of chemosensory experience. **A**. Worm mutants were exposed to BA 1/300 for 60 hours post-hatch in order to generate an olfactory imprinting. This odor-exposure was reiterated for several consecutive generations. To assess stable reshaping of the BA-chemoattraction, the F1 to F5 progeny issued from every BA-exposed generation (and from unexposed as control) were grown without BA exposure for 5 generations. Mean migration indices of the unexposed progeny - from F1 up to F5 - were determined and compared to the mean migration indices of naïve animals of the same genotype, as described. Each chemotaxis assay included 20 worms and was repeated four times. **B**. Consecutive generations of *ife-4 (ok320)* worms were exposed (or unexposed for naïve controls) to BA1/300 during 60, 36 or 12 hours post-hatch. To assess stable reshaping of the BA-chemoattraction, the F1 to F5 progeny issued from the three imprinting intervals was assessed. Each chemotaxis assay included 20 worms and was repeated four times.

## Discussion

We pinpointed several molecular mechanisms that control the temporal settings of chemosensory adaptation in *C. elegans* individuals and populations. Our data suggest that temporal shifts in neuronal maturation of individuals by epigenetic mechanisms could dramatically influence how and how fast experience can be cumulated, accounted and integrated by animal populations.

Chemo-attraction implementation progresses throughout wild-type worms development, despite a latency period concomitant with the olfactory plasticity period. Worms lacking functional *hbl-1, hda-3, spr-1 and spr-5* genes express chemo-attraction precociously without this latency period. Such precocity suppressed olfactory imprinting and modified the processing of early olfactory experience. This modified processing was previously observed in the imprinting mutants *sra-11* and *ttx-3* (5), but the mechanisms implicated in this phenomenon remain unclear. In contrast, worms in which *hbl-1* was specifically overexpressed in AIY interneuron and worms in which *hbl-1* expression was maintained overtime due to the lack of the putative inhibitor IFE-4 displayed a lengthened latency period of plasticity, up to adulthood. This could be explained by the maintenance of AIY in an immature highly plastic state by the continuous expression of the HBL-1 timer.

The timing of chemosensory adaptation we describe here displays some analogy with the timing of larval development in *C. elegans* (10, 11). Indeed, in both cases, loss of function or persistent expression of timers blocks the right progression of the temporal series. In the developmental heterochronic pathway, loss of function mutations in the *lin-14, lin-28* or *lin-41* genes result in skipping the L1, L2 or L3 larval stages, respectively, and promote premature expression of the next stage Instead, gain of function mutations of these genes produce overtime reiteration of the same respective stages. In the behavioural heterochrony pathway, loss of function in the *hbl-1, spr-1, spr-5* and *hda-3* genes induce a premature expression of the innate chemosensory responses, skipping the imprinting plasticity period, while HBL-1 gain of function in *ife-4* and *ttx3::hbl-1* mutants maintained the ability to imprint odours over life-time. These results seem to indicate that any behaviour open to experience-induced adaptation will not be expressed while behavioural adaptation can happen.

LSD1/HDAC1/2/CoREST (LHC) complexes modulate gene expression through interaction with different zinc-finger transcription factors (30), including REST/NRSF and Tramtrack88 in mammalian and fly neurons, respectively (19). Besides SPR-5/LSD1, SPR-1/CoREST may bind members of the HDAC family through its two SANT domains (23). *spr-1* and *hda-3* mutants display a precocious expression of the innate chemo-attraction, suggesting SPR-1 could associate with HDA-3 to control AIY plasticity timing.

Our results support the existence of a worm LHC complex comprised of SPR-1/CoREST, SPR-5/LSD1 and HDA-3/HDAC1. In the absence of a REST-like protein it may associate with HBL-1 to regulate gene expression in *C. elegans* neurons. Interestingly in a study focused on the epigenetic control of olfactory receptor (OR) choice by mouse olfactory neurons (ORN), it has been shown that the role of LSD-1 is to remove the repressive methylation me2 and me3 marks on the lysine 9 of H3 histones (31). The down-regulation of LSD-1 as ORN mature timely links OR gene expression to neuronal differentiation in the mouse. Our results suggest that SPR-5/LSD-1 may also control OR genes expression in the worm; the absence of OR repression in spr-5 mutants would impairs chemo-sensory neuron plasticity.

Demethylation by LSD1 and deacetylation by HDAC result in chromatin compaction and prevent transcription factors, regulatory complexes and RNA polymerases to bind DNA, all well known epigenetic mechanisms. Interestingly, LSD1 is also an integral component of the Mi-2/nucleosome remodelling and deacetylase (NuRD) complex (32), which mediate transcription repression by distinct factors including Hunchback and Ikaros (33). Noteworthy, Ikaros, Hunchback and Tramtrack88 also recruit LSD1 (34).

We report here opposite roles for the *C. elegans* histone acetylases HDA-2 and HDA-3 in the regulation of the chemosensory behaviour adaptation. Based on primary sequences comparison (data not shown), the worm HDA-3 and HDA-2 could be the respective orthologs of the mammalian HDAC1 and HDAC2. Strikingly, HDAC2, but not HDAC1, negatively regulates learning and memory in the mouse. HDAC-2 acts as a repressor of synaptic remodelling and plasticity genes (35) and HDAC2 inhibition restores cognitive abilities in a mouse model of Alzheimer’s disease (36). Our findings suggest the opposite functions for HDA-3/HDAC1 and HDA-2/HDAC2 in neuronal plasticity and maturation might be evolutionary conserved. A putative HBL-1/LHC complex would thus epigenetically control the timing of the plasticity periods within AIY interneurons. Furthermore, IFE-4 seems to be essential to inhibit HBL-1 translation.

Our results suggest that worms produce heritable olfactory imprints exclusively during the AIY plasticity period, while the innate chemo-attraction is repressed. In multigenerational imprinting situation, odour-exposed worms would add imprints they produce to imprints inherited from past generations, leading to imprint accumulation throughout generations, reaching an effective threshold that triggers integration and stable inheritance of the newly acquired experience.

In this paper, we tried to unveil some of the relationships between the different temporal variables that may determine the rate at which multigenerational experience stably change animal behaviours. As schematically outlined in Figure 6, it could be inferred from our data that 1) the onset of an adult behaviour is inhibited while the period of plasticity is maintained; 2) the period of plasticity delimits the amount of individual acquired experience imprinting the adult behaviour, and; 3) the amount of acquired experience constraints the adaptation of the populations. Specific genetic and epigenetic factors should time the functional maturation of neurons responsible for a given behaviour. Then the time-length of juvenile periods of behavioural plasticity would in turn determine the amount of experience that can be acquired during a generation. Stable changes in population behaviour would be acquired through the integration of cumulated heritable experience over multiple generations. The rate of stable behavioural adaptation would thus be determined by the amount of experience acquired by each generation. In brief, neuronal maturation speed is expected to be responsible for the rate innate behaviour change, according to ancestral experience.

**Figure 6.**
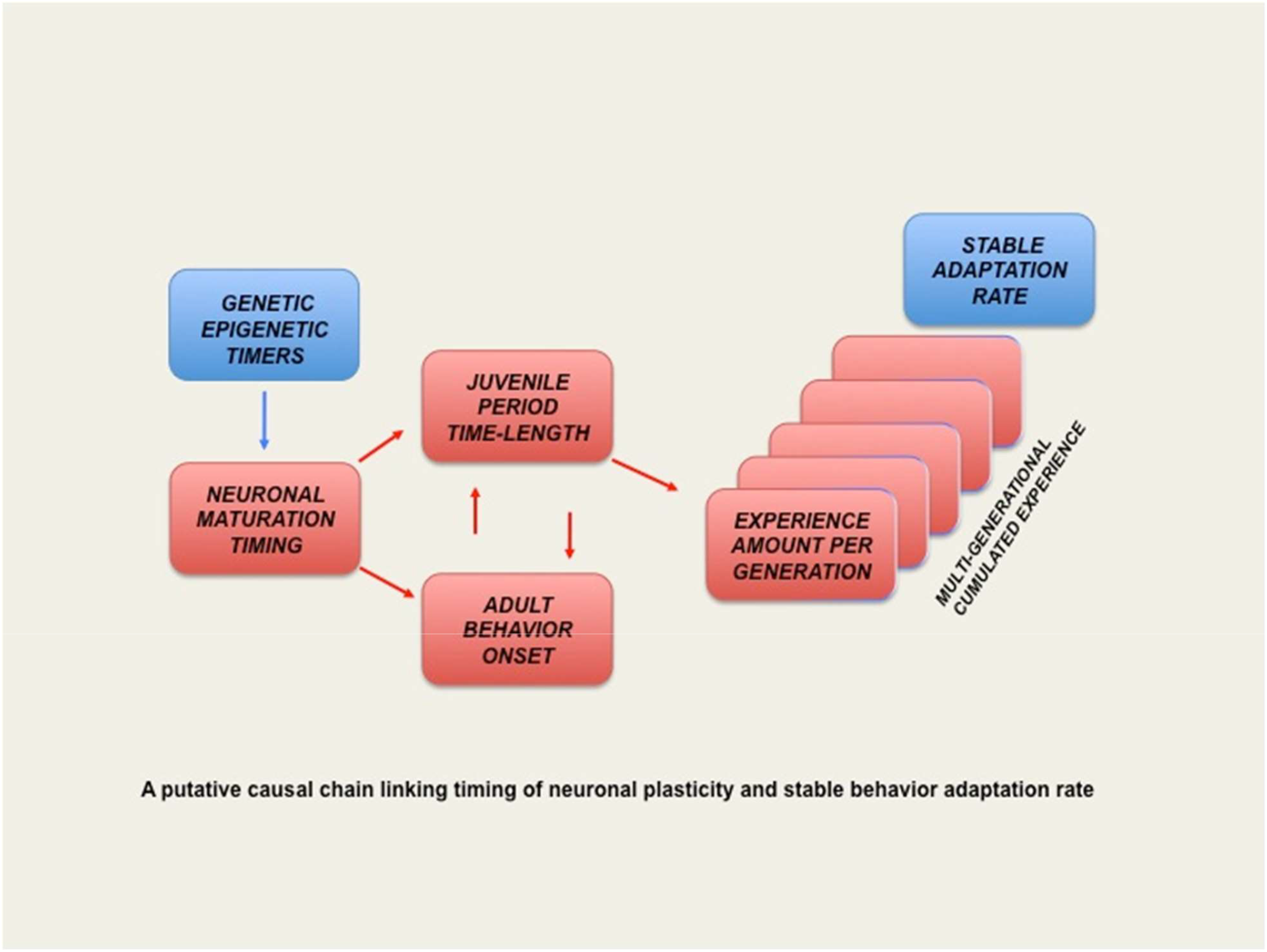
A putative causal chain linking the timing of neuronal plasticity to the stable behaviour adaptation rate.

Through yet unknown mechanisms, an environmentally induced reversible adaptation may evolve towards stable expression without triggering environmental stimuli (29). In particular, it is now well accepted that epigenetic alterations may contribute to establish phenotypic divergence among genetically identical animals exposed to different experiences (37). Other works also indicate that temporal shifts of gene expression during development have been recognized as potential drivers of speciation (11, 38, 39). Evidence also indicates that learning and behavioural/cultural shifts could support fast reproductive isolation (40, 41). The present work suggests that temporal shifts in epigenetically controlled neuronal maturation could entail dramatic consequences on stable behavioural adaptations.

## Material and Methods

*C. elegans* strains were maintained at 20°C under standard growth conditions (Brenner, 1974). Synchronized populations of worms were used to perform all experiments. All confocal images were acquired using a Zeiss LSM780 microscope. The HBL-1 antibodies were custom-made. Supplementary Materials and Methods contain detailed descriptions on the worm strains, and on imaging, Western blotting and chemotaxis assay protocols. Chemotaxis assay is schematized in the Supplemental Figure 3.

We have no competing interests.

## Footnotes

### Author contributions

MG carried out the molecular lab work, participated in data analysis, FF participated in the design of the study; DAFdA edited the manuscript; AG, KB and YC carried out molecular lab work and behavioural experiments; MPB collected microscopy data; JJR conceived and coordinated the study, performed experiments, carried out the statistical analyses and draft the manuscript. All authors gave final approval for publication.

## Acknowledgements

We thank O. Hobert, D. Baillie, M. Delattre and I. Greenwald for worm strains and plasmids. Other strains used in this study were provided by the Caenorhabditis Genetics Center, which is funded by the National Institutes of Health (NIH) Office of Research Infrastructure Programs (P40 OD010440). This work was supported by a grant of the “Agence Nationale de la Recherche” ANR-12-Bioadapt-0022.

## Notes

### Competing Interest Statement

The authors have declared no competing interest.

