## Supplemental text and supplemental Figures for "Epigenetic regulation clocks the multigenerational olfactory imprinting in *C. elegans*"

### Supplemental Material and Methods

#### *Worm Strains*

*C. elegans* strains were maintained at 20°C under standard growth conditions (Brenner, 1974). Strains used in this study include: N2, *hbl-1 (ve18)*, *hbl-1 (mg285)*, *ife-4(ok320)*, *spr-1(ar200)*, *spr-1(ar205)*, *spr-5(by119)*, *spr-5*(*ar197), spr-5*(*by101) hda-2(ok1479)*, *hda-3(ok1991)* and *hda-4(ok518).* The reporter strains analyzed were: BW1932 (hbl-1::gfp-hbl-1 reporter) and BW1981 (hbl-1::gfp-unc-54 reporter (Fay et al, 1999) and the *ott4607* strain expressing CherryFP under the *ttx-3* promoter B (gift of O. Hobert). The *sEx15051* strain contained the extrachromosomal *ife-4*::*GFP* reporter array (gift of D. Baillie). N2 worms carrying the [*pttx-3::hbl-1;sca- 1::gfp*] transgene were obtained as follows: the *hbl-1* gene was amplified by PCRfrom *C.elegans* genomic DNA and subcloned in pCR-XL-TOPO®. A 4060bp Kpn- I/Not-I fragment was then combined in a ligation reaction with a Kpn-I/Not-I fragment of a vector containing the 243bp AIY-regulatory element of the *ttx-3* locus (gift from O. Hobert, Columbia University, New York). The *ttx-3::hbl-1* vector was injected into N2 wild-type animals at 15 ng/µl using pGK10 as a co-injection marker.

#### *Imaging*

Worms were immobilized with 50 µM Na-azide in M9 buffer. All confocal microscope images were acquired using a Zeiss LSM780 microscope with a 63× NA 1.4 oil immersion objective. Image stacks were captured and maximum intensity projections were obtained using ZEN 2010 software.

***Western Blotting***

Worms were washed in M9 buffer and spun down at 6000 rpm for 30 seconds. An equal volume of lysis buffer containing 50 mM Tris pH7.5, 50mM NaCl and fresh protease inhibitors: (aprotinin, pepstatinA, leupeptin, PMSF) was then added. After freezing in liquid nitrogen, the solution was quickly thawed in a water bath and sonicated on ice (3x 15s). Total protein extracts **(**100 µg) were used for SDS-Page/Western blots. Two antigenic peptides of HBL-1 (aminoacid 1-15 and 863-877) were designed and used to immunize rabbits. HBL-1 antibodies were produced using the 28-day Polyclonal Super Speedy protocol (Eurogentec). Western blot were scanned, quantified by ImageJ and the intensity of HBL-1 bands were normalized to those of actin (mouse monoclonal antibodies clone C4; Millipore).

***Chemotaxis assays and odor-exposure***

Synchronized populations were obtained by allowing about ten adult worms to spawn embryos during one hour on 6 cm OP50 loaded NGM plates.

In order to induce the olfactory imprint, a 4µl drop of 1/300 diluted benzaldehyde was suspended on the lids of worm culture plates during the indicated periods of time at the indicated development stages.

Population chemotaxis assays were performed at the indicated ages on 12 cm squared low-salt agar plates. The assay used in this study is based on the population chemotaxis assay originally described (Bargmann, C.I. Chemosensation in *C. elegans* (October 25, 2006), *WormBook*, ed. The *C. elegans* Research Community, WormBook, doi/10.1895/wormbook.1.123.1). Worms are placed in the center of low-salt (1mM Ca, 1mM Mg, 5mM KPO4) agar plates and allowed crawl up a gradient of an attractive odorant during approximately one hour. To form odor gradients odors are spotted at one end of the plate, while the same volume of water is spotted at the opposite end. 1 µl of sodium azide (NaN3) is added in the agar at both ends of the plate to immobilize animals that reached the odor or the water positions.

Compared to the original assay, several modifications were made in the procedure and in the way chemotaxis indices were calculated. Changes aimed at more accurately compare the chemo-attraction of worm populations to moderately attractive odorants (as the 1/300 dilution of benzaldehyde used here), using a lower number of animals (20 worms/assay) (Supplementary Figure 3).

**Procedure**: Agar plates were dried at room temperature at last during three days prior to the assays. Nutrition state, humidity, temperature shifts, or worm-to-worm interactions could interfere with chemotaxis, especially when worms migrated up gradients of mildly attractive odorants. In order to eliminate these potential sources of variability, 20 worms were individually transferred from the culture dish to the middle line of the assay plate. To establish a homogeneous odor gradient, so that all worms were submitted to the same olfactory stimulus, 3 drops of odor-dilution (4 µl each) were suspended on the lids at one side on the squared plate, each placed at a distance of 3 cm from the others.

**Mean migration indices**: In the original population assay, migration to an odor source was translated into a chemotaxis index. Index was calculated at the steady state as a ratio, i.e. the proportion of worms that reached the odor position minus the proportion that reached the diluent position divided by the total number of worms in the assay. However, worms in a gradient of mildly attractive odors (the case here) did not move straight forward. They often migrated up and down the gradient, so during the time of the assay, most worms sat at different places between the starting position and the odor source. The way chemotaxis index was calculated so as to sensitize the assay by taking into account all worm moves during the course of the assay, i.e. before the steady state. In this study, larval and adult stages were assayed. Larvae migrated towards odor sources more slowly than adults and never reached the steady state within one hour. Squared plates facilitate the indexation of every worm position between the starting line (time 0, position 0 cm) and the odor source (position + 6 cm), usually four times at 10, 20, 30 and 40 minutes from time 0. Worms that migrated down the odor gradients were negatively indexed (from 0 to – 6). The data collected in that way could be seen as a motion picture of all worm moves, with 4 images per hour. The mean value of all indexed positions (in cm from the starting line) of each of the 20 worms involved in the assay represented the mean migration index.

Chemotaxis assays were always performed so as to compare worm populations. Populations to be compared were synchronized and always blindly tested the same day using the same batch of assay plates.

**Imprinting indices:** Imprinting indices were calculated based on the difference between the mean migration index of 20 naive and the mean migration index of 20 odor-exposed worms, both of the same genotype and age.

**Statistical analysis**

Mean migration and imprinting indices (+/- s.e.) were compared using unpaired data with unequal variance Student t tests (KaleidaGraph).

### Supplemental Figures legends

#### Supplemental Figure 1

**A. The *hbl-1::GFP-hbl-1* reporter is expressed in the AIY interneurons during all developmental stages.**

Confocal micrographs show the co-localization of *ttx-3::mCherryFP* (*ott4607*, red fluorescence) and *hbl-1::gfp-hbl-1* (BW1932, green fluorescence) reporters in L1 (A, B, C), L2 (D, E, F), L3 (G, H, I) larvae and in adult worms (J, K, L). Merged images are shown in the right column. All figures are oriented with anterior to the left and posterior to the right. The scale bar indicates 10 µm.

**B. Expression of an *ife-4::GFP* reporter in the AIY interneurons starts after 12 hours post-hatch.**

Merged images of confocal micrographs are shown in the right column. *ife-4::GFP* fluorescence is not observed in AIY during the L1 stage (A, B, C). *ife-4::GFP* (green fluorescence) begins during the L2 stage (D, E, F) and persists through the L3 (G, H, I) and the L4 (J, K, L) stages. *ttx-3::mCherryFP* (*ott4607,* red fluorescence) co-localised with *ife-4::GFP*. All figures are oriented with anterior to the left and posterior to the right. The scale bar indicates 10 µm.

#### Supplemental Figure 2

**IFE-4 affects the *hbl-1* translational efficiency during larval development.**

Total protein extracts were collected from synchronized populations of wild-type N2 and *hbl-1(mg285)* and *ife-4(ok320)* mutants of the indicated ages. Total protein extracts were then migrated in SDS-polyacrylamide gels. HBL-1 was detected by Western blot using an anti-HBL-1 antiserum. The intensity of HBL-1 bands was normalized to the bands of actin (% hbl-1 expression).

#### Supplemental Figure 3

**Schematisation of the chemotaxis assay.** 20 worms are placed at the starting line of the plate at time 0. The odour and the water are placed at opposite extremities of the plate’s lid. Following a pre-determined schedule, the free progression of the worms on the agar is recorded. The mean migration index of the assay is given by the equation.

Supplemental Figure 1A

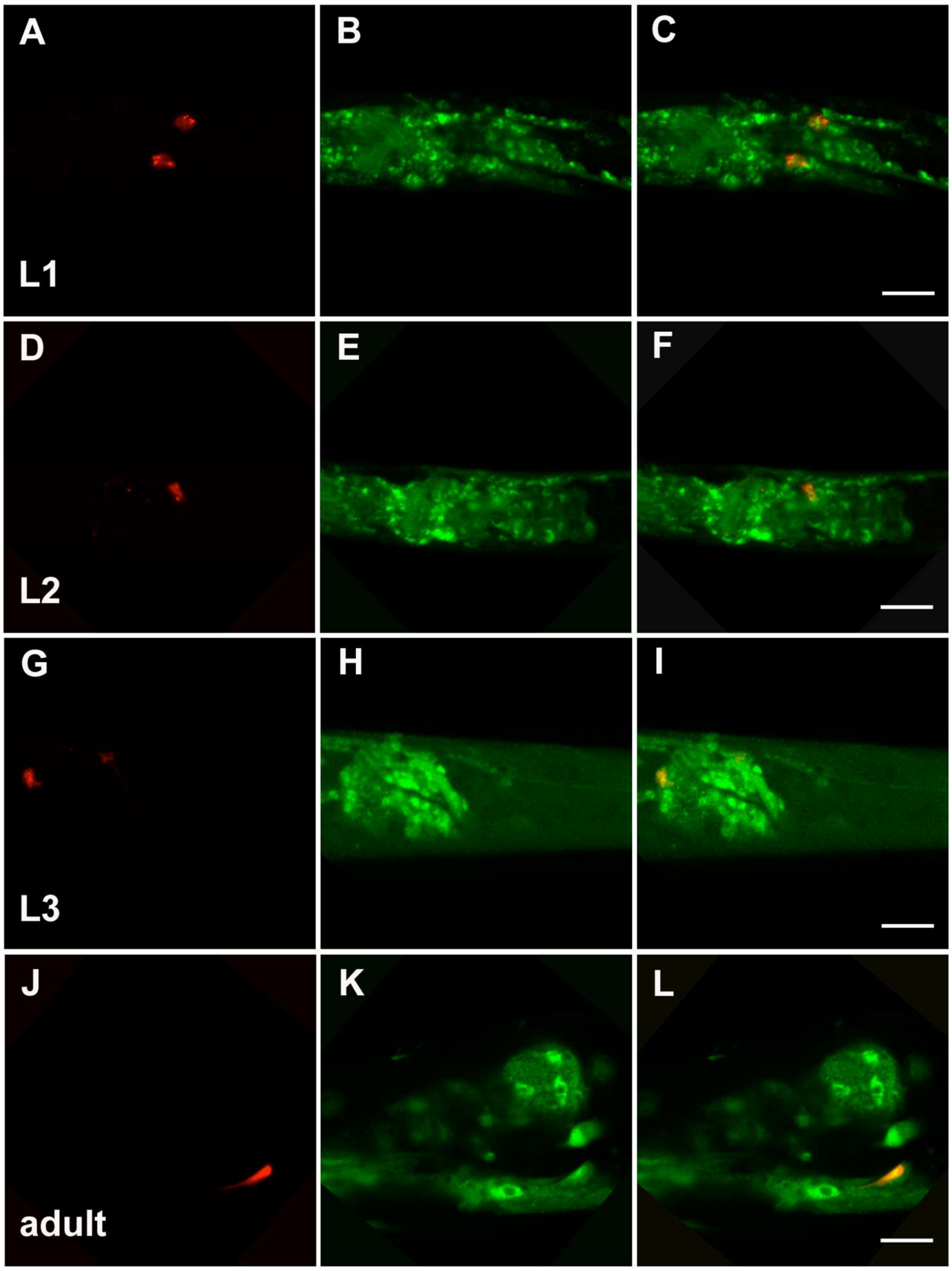

**Supplemental Figure 1B**

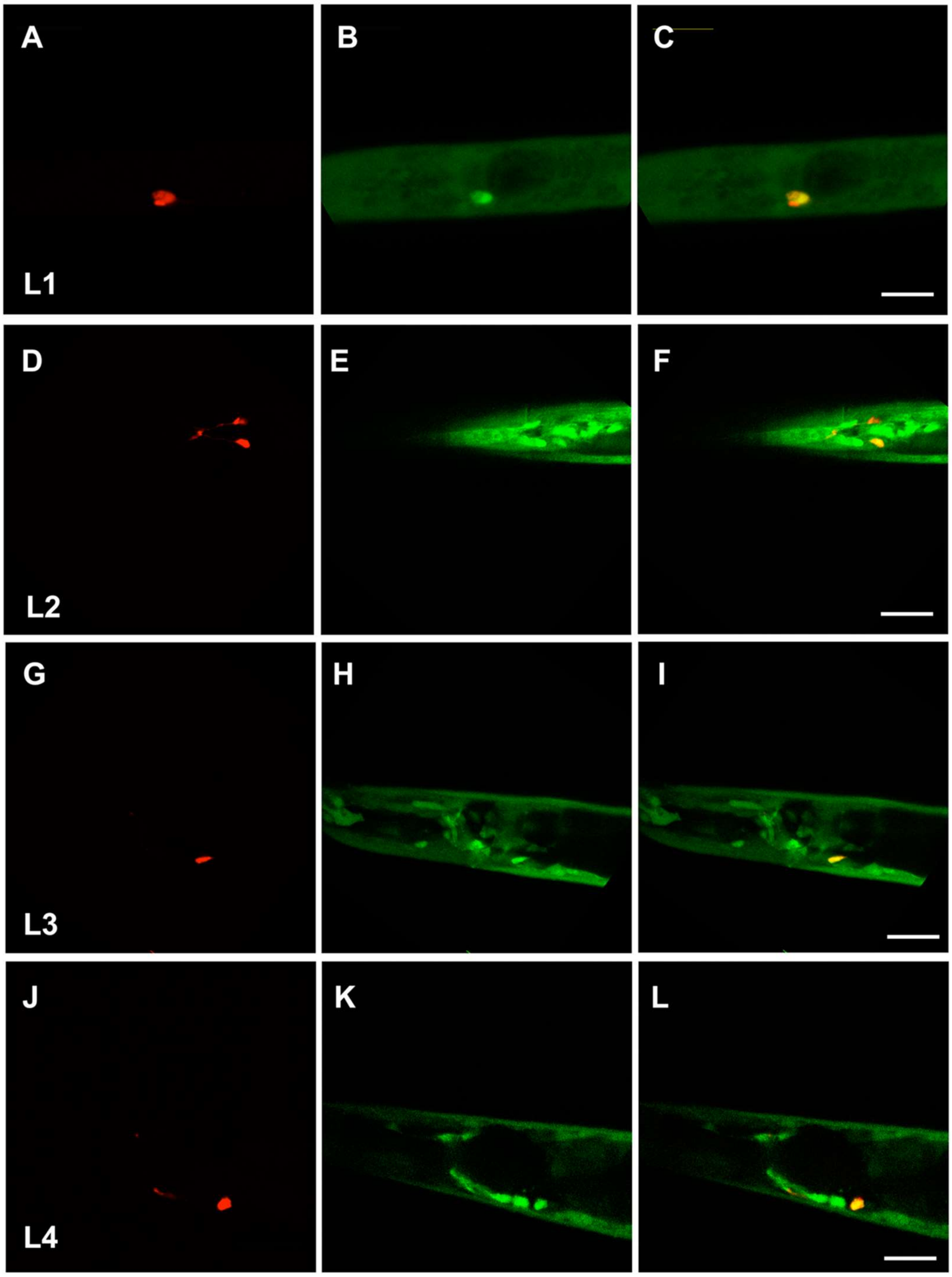

**Supplemental Figure 2**

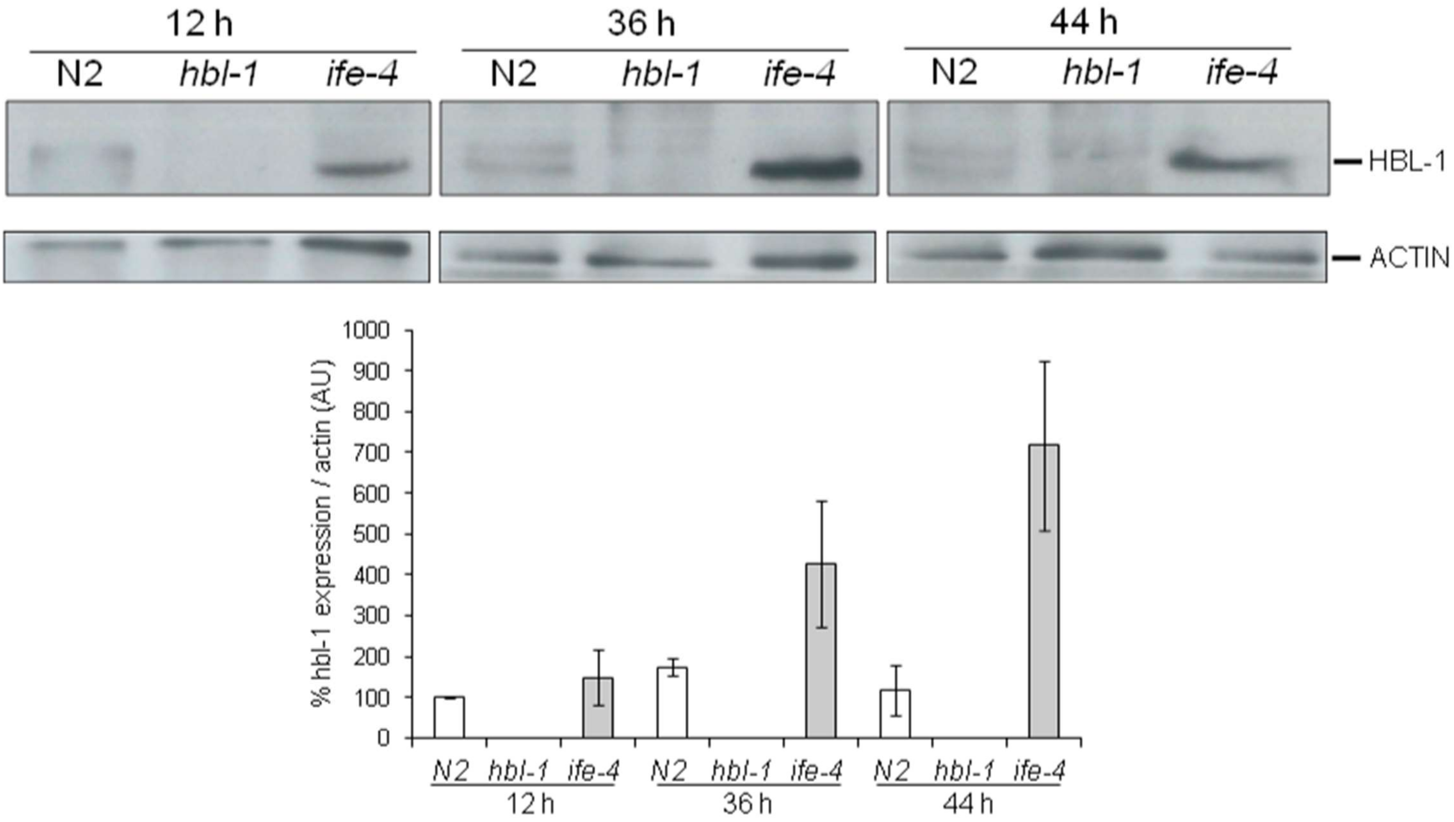

**Supplemental Figure 3**

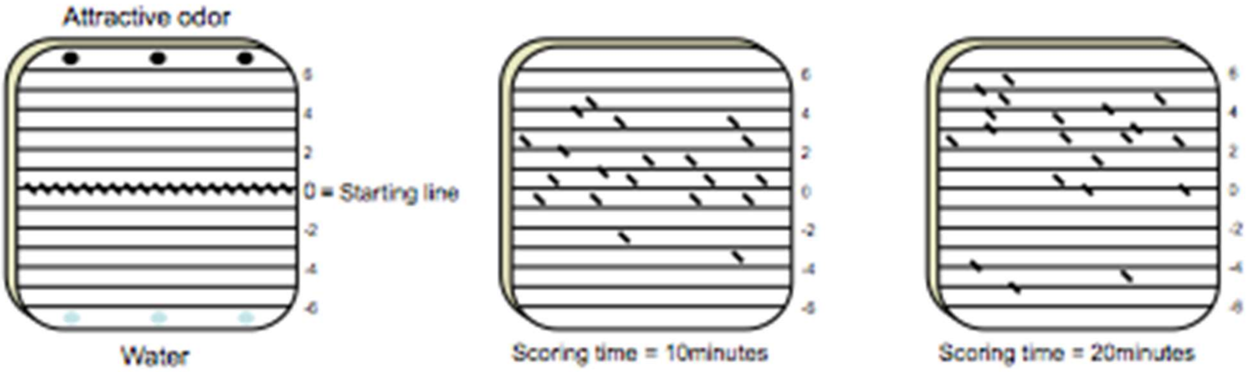


$$MMI= \frac{1}{m}\sum_{i=n}^{m}\chi i$$
